# Drug Response Prediction Provides a Biologically Relevant Benchmark for Perturbation Response Models

**DOI:** 10.64898/2025.12.09.693213

**Authors:** Niek Brouwer, Martin Damyanov, Jonas Argelo, Daniël J. Vis, Marcel J. T. Reinders, Lodewyk F. A. Wessels

**Affiliations:** Division of Molecular Carcinogenesis, Oncode Institute, The Netherlands Cancer Institute, 1066 CX Amsterdam, The Netherlands; Department of Electrical Engineering, Mathematics and Computer Science, Delft University of Technology, 2628 XE Delft, The Netherlands

## Abstract

Perturbation response models (PRMs) predict transcriptional responses to interventions such as gene knockout or drug treatments. While simple baseline models match PRMs in reconstructing post-treatment profiles, we show that drug response predictors trained on PRM-generated profiles significantly outperform those trained on baseline or pre-treatment profiles. Unlike simple baseline models, PRMs preserve variation in response-relevant genes, making them a highly valuable tool for predicting drug response.

---

Identifying effective cancer treatments for individual tumors remains a central challenge in oncology. Predictive models that estimate drug sensitivity from molecular data hold promise to guide treatment selection. Yet, current approaches based on pre-treatment gene expression profiles often fail to generalize across cell lines or tumor types (Adam et al. [2020], Partin et al. [2025]). Limited performance arises because sensitivity and resistance are driven by dynamic transcriptional programs that are not observable before treatment.

In contrast, post-treatment gene expression profiles capture the drug-induced changes underlying response phenotypes (McFarland et al. [2020], Srivatsan et al. [2020]). Recently proposed perturbation response models (PRMs), such as CPA (Lotfollahi et al. [2023]), GEARS (Roohani et al. [2023]), and scFoundation (Hao et al. [2024]), can predict post-treatment profiles from pre-treatment data, enabling modeling of cellular responses without additional experiments.

PRMs have so far been evaluated mainly on their ability to reconstruct measured post-treatment profiles, directly comparing measured and predicted gene expression values. Using reconstruction accuracy as a measure, deep learning-based PRMs fail to outperform simple baseline models that predict no or average effects (Ahlmann-Eltze et al. [2025]). However, to comprehensively evaluate and interpret the effects of perturbations, it is essential to establish connections between predicted profiles and phenotypic outcomes. Therefore, PRMs should also be assessed on their capacity to capture biologically meaningful information relevant for predicting and understanding treatment outcomes.

In this study, we establish whether drug response predictors trained on post-treatment profiles predicted by PRMs outperform those trained on pre-treatment profiles, and, importantly, whether they also outperform predictors trained on profiles generated by simple baseline models (Fig. 1a).

**Figure 1:**
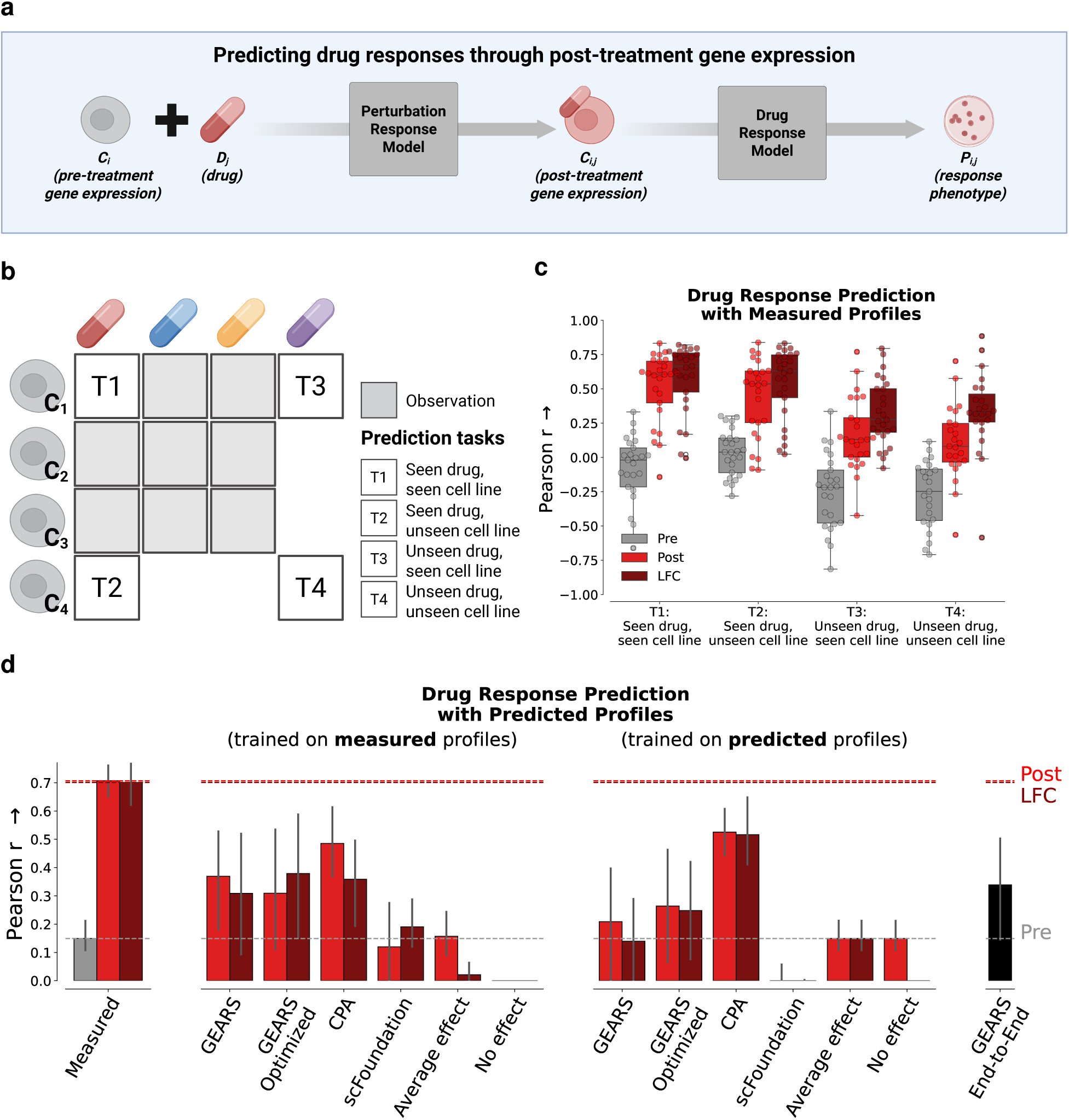
Drug response prediction with predicted post-treatment gene expression profiles. **a,** Overview of the prediction pipeline. First, post-treatment profiles are predicted with a perturbation response model. Next, drug response models are trained on these profiles to predict drug response phenotypes. **b,** Overview of different prediction tasks. T1: Unseen (cell line, drug) pair, T2: unseen cell line, T3: unseen drug, T4: Unseen cell line and unseen drug. **c,** Performance of drug response models trained and tested on measured pre- post-, and LFC gene expression profiles from the McFarland dataset. To mitigate confounding by tissue lineage, cell lines from the same tissue type were assigned exclusively to either the training or test set. Performance was evaluated by grouping predictions by drug and tissue type, and computing the Pearson correlation coefficient between predicted and observed AUDRC values. **d,** Performance of drug response models trained and tested on measured profiles; trained on measured profiles and tested on predicted profiles; and trained and tested on predicted profiles.

First, we assessed the potential of post-treatment gene expression as informative input to drug response predictors. To this end, we determined how well we could predict drug sensitivity, using a dataset of experimentally measured post-treatment gene expression and matching response measurements (McFarland et al. [2020]). The McFarland dataset comprises 170 cell lines from 14 tissue types, each treated with nine drugs. For all (cell line, drug) pairs, pre- and post-treatment gene expression profiles were available. Additionally, we computed the log fold change (LFC) between pre- and post-treatment profiles. Each representation was used as input to regularized linear models predicting drug sensitivity as quantified by 1 - the observed Area Under the Dose–Response Curve (1-AUDRC) (Methods).

We defined different prediction tasks that differed in whether cell lines and drugs in the test set had been observed during training (Fig. 1b, Extended Data Fig. 1). Models trained on post-treatment gene expression, particularly LFC, consistently improved performance across all tasks, compared to pre-treatment data. Notably, performance did not decline when cell lines were unseen, indicating that the models captured molecular features of drug sensitivity that generalize across tissue types. In contrast, performance dropped substantially when the drug was unseen, demonstrating that predicting responses to entirely novel drugs remains more challenging (Fig. 1c).

After establishing that post-treatment profiles provide a more informative basis for drug response prediction than pre-treatment profiles, we used three PRMs (CPA (Lotfollahi et al. [2023]), GEARS (Roohani et al. [2023]), and scFoundation (Hao et al. [2024])), trained on the Sciplex3 dataset (Srivatsan et al. [2020]), to generate post-treatment gene expression profiles (Methods).

In addition to the original model implementations, we developed two GEARS variants to assess model behavior further. First, we generated “GEARS optimized” by removing excessive regularization, which improved performance on a genetic perturbation screen (Norman et al. [2019]) (Extended Data Fig. 2). Second, we implemented “GEARS End-to-End”, an end-to-end variant that directly predicted AUDRC values from latent embeddings rather than explicitly generating post-treatment gene expression profiles (Methods).

Lastly, we also included two baseline models in the comparison: a “No effect” model that predicts no change in gene expression upon perturbation, and an “Average effect” model that always predicts the average measured post-treatment profile. Consistent with earlier studies (Ahlmann-Eltze et al. [2025], Bendidi et al. [2024], Li et al. [2024]), we found that the reconstruction accuracy of profiles predicted by PRMs was comparable to that of these baselines in terms of Mean Squared Error (MSE) and Pearson correlation coefficient (PCC) (Extended Data Fig. 3).

Next, we evaluated whether profiles predicted by PRMs and baseline models were also equally informative for predicting drug responses. To this end, we trained regularized linear regressors on either measured or predicted post-treatment gene expression profiles to predict 1-AUDRC values (Methods).

Remarkably, drug response models trained on measured profiles predicted drug sensitivity more accurately from profiles generated by CPA and GEARS than from those generated by baseline models (Fig. 1d, Panel 2; Extended Fig. 4). Moreover, training the model on CPA-predicted profiles further improved accuracy (Fig. 1d, Panel 3), suggesting that the predicted features form a denoised or stabilized representation of the biological signal. This interpretation was further supported by the model’s strong generalization when tested on measured profiles (Extended Data Fig. 5).

In contrast, the model trained on GEARS-predicted profiles showed lower accuracy than when trained on measured profiles (Fig. 1d, Panel 3), suggesting that although these profiles capture some biologically meaningful signals, the feature space is noisy or internally inconsistent. The optimized GEARS variant exhibited behavior comparable to that of the original GEARS model. Predictions based on scFoundation-predicted profiles consistently performed poorly, indicating that this model fails to recover response-associated gene expression changes.

Finally, GEARS End-to-End achieved accuracy equivalent to models that first predicted post-treatment profiles and then predicted drug response (Fig. 1d, Panel 4), suggesting that explicitly predicted gene expression profiles are potentially more informative for drug response prediction than latent embeddings.

All observed trends were consistent across multiple performance metrics, including the MSE between predicted and observed AUDRC values and top-10 retrieval accuracy (Extended Data Fig. 6).

Because baseline models generate identical profiles for all drugs, the additional variation introduced by PRMs likely drives their improved performance in predicting drug responses. To quantify this variation, we compared gene-gene correlations in measured and predicted post-treatment profiles. When computed across all cell lines, correlations were dominated by cell line-specific expression and were similar for measured profiles and for profiles predicted by PRMs and baseline models (Fig. 2a). In contrast, when computed within each cell line, correlations were preserved only in CPA-predicted profiles, indicating that CPA more accurately reconstructed the joint behavior of genes (Fig. 2b). GEARS-predicted profiles preserved correlations only weakly and in only one of the three cell lines, consistent with its variable performance (Extended Data Fig. 7).

**Figure 2:**
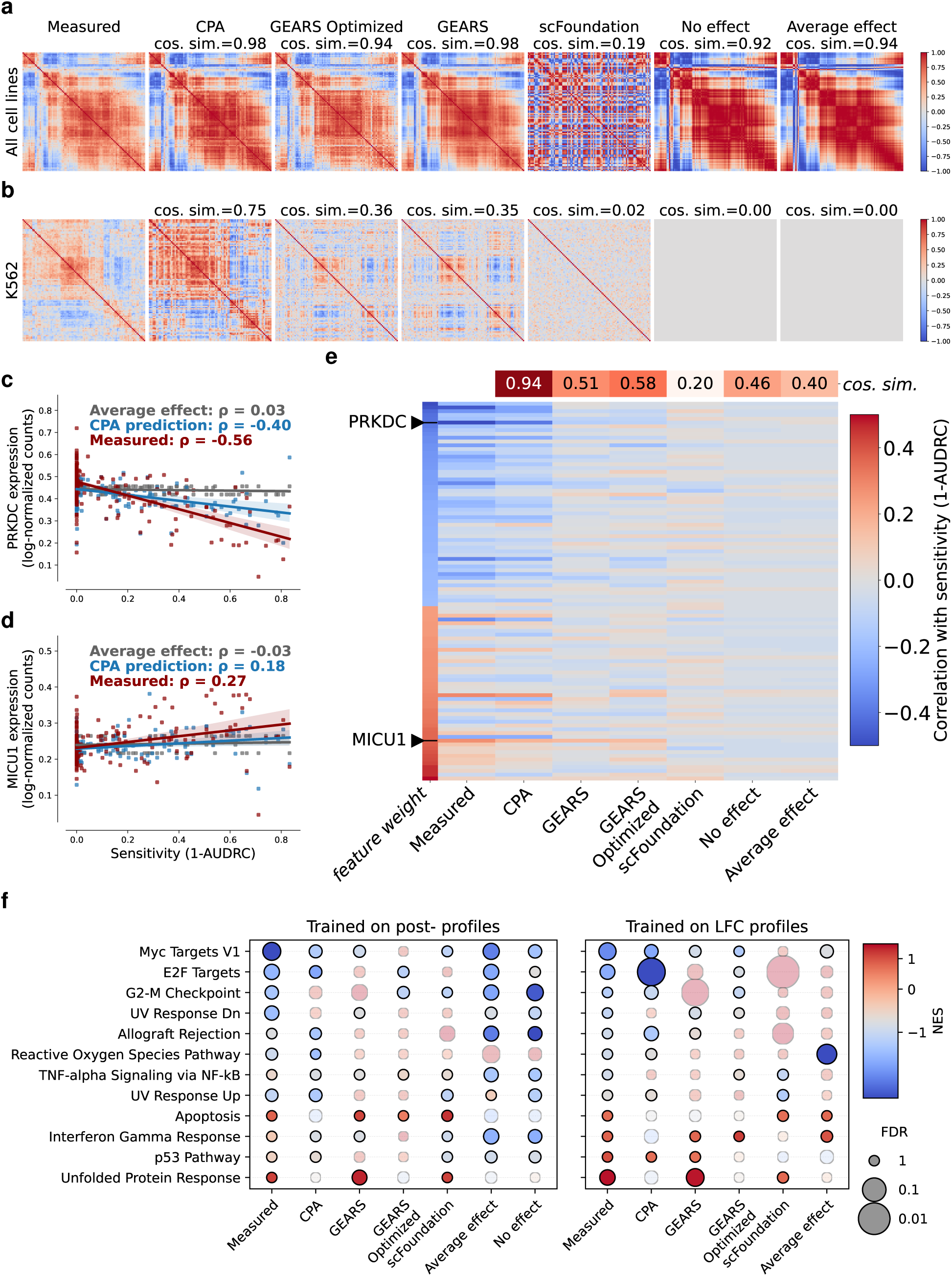
Identification of informative features for drug response prediction. **a,** Pairwise correlations between genes selected by drug response models trained on measured profiles. Correlations are computed across cell lines. **b,** Gene-gene correlations computed for K562 cell line. **c,** Gene expression of PRKDC (part of the G2/M checkpoint gene set) across (cell line, drug) pairs against drug sensitivity. Gene expression is downregulated for more sensitive pairs. The correlation is conserved in profiles predicted by CPA. **d,** Gene expression of MICU1 (part of the Reactive Oxygen Species pathway gene set) across (cell line drug) pairs against drug sensitivity. Gene expression is upregulated for more sensitive pairs. The correlation is conserved in profiles predicted by CPA. **e,** Correlations between expression values and drug sensitivity for the genes contributing most to the drug response model trained on measured profiles. As expected, correlations are strong for measured profiles, but correlations are also largely preserved in the profiles predicted by CPA and to some extent by GEARS. **f,** Gene set enrichment for genes selected by drug response models trained on measured and predicted post-treatment gene expression profiles. Dots are colored by Normalized Enrichment Score (NES), sized by adjusted p-value (FDR), and faded if the sign of the NES does not match that of the model trained on measured profiles.

To determine whether this variation was linked to drug sensitivity, we computed correlations between gene expression and 1-AUDRC values. Measured expression values correlated with sensitivity as expected: genes which were assigned negative feature weights by the linear regressors showed lower expression in sensitive cell lines (Fig. 2c), whereas genes with positive weights showed higher expression (Fig. 2d). These correlations were well preserved in profiles predicted by CPA (cosine similarity = 0.94), and to a lesser extent by GEARS (cosine similarity = 0.51 and 0.58), but were largely absent in predictions from scFoundation (cosine similarity = 0.20) and baseline models (cosine similarity = 0.46 and 0.40) (Fig. 2e).

Finally, we assessed whether the genes selected by drug response models corresponded to common biological programs by performing gene set enrichment analysis (GSEA) on the feature weights (Methods). This analysis identified Hallmark gene sets (Liberzon et al. [2015]) whose expression was consistently associated with sensitivity (Fig. 2f).

Models trained on measured profiles selected features enriched for apoptosis and stress-related programs, such as the Unfolded Protein Response and p53 Pathway. These positively enriched pathways comprised genes that consistently received positive weights, indicating that higher expression of these genes was associated with increased drug sensitivity. In contrast, cell cycle-related pathways, such as the G2/M checkpoint, E2F targets, and MYC targets, showed strong negative enrichment, indicating that genes in these pathways were assigned negative weights and that higher expression levels were inversely associated with drug sensitivity. These enrichment patterns closely resembled those obtained from drug response models trained on measured profiles in the McFarland dataset, indicating that the selected features reflect robust molecular mechanisms associated with treatment response (Extended Data Fig. 8). Models trained on CPA-predicted profiles, which achieved high predictive accuracy, displayed similar enrichment for E2F Targets, MYC Targets, G2/M Checkpoint, and Allograft Rejection, but lacked positive enrichment for stress-related pathways.

Taken together, these findings show that PRM-predicted profiles carry a biologically meaningful signal. These signals are not reflected in reconstruction accuracy metrics, such as the MSE and PCC, suggesting that these metrics are not the most relevant for benchmarking performance. Although alternative evaluation strategies are emerging (Viñas Torné et al. [2025], Miller et al.), evaluating PRMs through challenging and biologically interesting downstream tasks provides more meaningful and practical insights.

Modeling drug response from predicted post-treatment profiles addresses the limited performance of approaches that rely solely on pre-treatment expression profiles (Adam et al. [2020], Partin et al. [2025]). Moreover, because PRMs can efficiently generate post-treatment profiles for different conditions, combining PRMs with drug response predictors is promising to enable rapid exploration of large intervention spaces.

Since predictive performance achieved with predicted profiles did not yet reach the level obtained with measured profiles, PRMs would benefit from incorporating prior information on gene function and from improving the modeling of relative expression changes. These directions will help position perturbation response modeling as a tool for addressing biological and clinical questions, rather than as an isolated methodological development.

## Online Methods

### Data Preprocessing

All gene expression matrices were unified to include human protein-coding genes and common mitochondrial genes, totaling 19.264 genes. If symbols were missing, they were imputed with zero values. We removed cells with fewer than 200 expressed genes. Subsequently, raw gene expression counts were normalized using total-count normalization to ensure comparability across cells, and log-transformation was applied using the natural logarithm plus one (ln1p) to stabilize variance and reduce the effect of outliers.

We computed pseudobulk gene expression estimates from single-cell data by averaging the processed (log-transformed) expression values across cells within each (cell line, drug) group to obtain condition- and cell-line-specific transcriptional profiles. Drug and cell line annotations were taken from the original studies accompanying the datasets. The mean log fold change (LFC) per gene was computed by subtracting the average log-transformed control expression from post-treatment expression within each cell line.

### Differential Expression Analysis

Differential gene expression between untreated and treated cell lines was assessed using the `rank_genes_groups` function from the scanpy package, comparing each drug-treated condition against a shared control group (DMSO treatment during the same time). All genes in the dataset were included in the ranking, and analysis was performed on the processed (non-raw) expression matrix. Genes were ranked by adjusted p-value.

### Drug Response Prediction with Measured Profiles

Prior to training, we filtered the data based on the coefficient of variation (CV) of AUDRC values to avoid predictions for tissue types and treatments with constant response values. For each (tissue, drug) group, the CV was computed as the standard deviation divided by the mean AUDRC. Groups with a CV below 0.5 were excluded from drug response prediction. To mitigate confounding by tissue lineage, cell lines from the same tissue type were assigned exclusively to either the training or test set. Finally, we applied naive feature selection based on variance filtering to reduce dimensionality and computational burden. We computed the variance of each gene across the untreated cells and selected the top 1000 genes with the highest variance.

Regularized linear regression models (ElasticNet) were trained with the 1-AUDRC value as the response variable. Hyperparameters (`alpha` and `l1_ratio`) were optimized using a 5-fold inner cross-validation grid search.

Trained models were evaluated on all (tissue, drug) groups with exhaustive leave-one-out cross-validation. Predictions for each (tissue, drug group) were evaluated using the Pearson correlation coefficient between the predicted and observed AUDRC values.

### Perturbation Response Models

All models were trained on a subset of the Sciplex3 dataset (Srivatsan et al. [2020]) that only comprised cells treated with drugs with known targets. To maximize on-target transcriptional effects, only treatments corresponding to the highest administered dosage per drug were retained.

GEARS and scFoundation use cell line–specific gene–gene co-expression graphs and are trained separately for each cell line, whereas CPA leverages larger training datasets across multiple cell lines. To generate predicted profiles for all (cell line, drug) pairs, each model was trained in a five-fold scheme, with 80% of the (cell line, drug) pairs used for training and the remaining 20% for inference.

### CPA

CPA (Lotfollahi et al. [2023]) was trained for 120 epochs following the procedures described in the original publication. The following hyper-parameters were used: learning rate = 0.003, autoencoder num layers = 4, autoencoder latent space dimension = 128, autoencoder layer size = 256, adversarial pretrain epochs = 25. adversarial num layers = 4, adversarial layer size = 64,

### GEARS

GEARS (Roohani et al. [2023]) was trained for 20 epochs following the procedure described in the original publication. As GEARS incorporates a knowledge graph of gene–gene relationships, drugs were represented by their target genes. The following hyperparameters were used: `hidden size = 64, learning rate = 0.001`. We trained the alternative version of GEARS with all regularization mechanisms removed: `gradient clipping = 0, weight decay = 0, and lr scheduler = none`.

### GEARS End-to-End

We modified the model’s forward pass to include an additional multi-layer perceptron (MLP) applied to the encoded representations, enabling direct prediction of drug sensitivity from pre-treatment gene expression profiles and perturbations. The embedding space retained the dimensionality of an expression profile, but components were not explicitly trained to represent genes. The regression head was trained using the mean squared error (MSE) loss between predicted and observed AUDRC values.

### scFoundation

scFoundation (Hao et al. [2024]) was trained for 20 epochs following the procedure described in the original publication. The following hyperparameters were used: `hidden_size = 512, learning_rate = 0.002, accumulation_steps = 1, highres = 0, finetune_method = frozen, and model_type = maeautobin.`

### Reconstruction Accuracy

To assess the reconstruction accuracy of predicted gene expression profiles, we computed the Mean Squared Error (MSE) and Pearson correlation coefficient (PCC) comparing predicted and measured values at the level of individual (cell line, drug) pairs. Predicted and measured gene expression profiles were averaged across replicates for each condition. Metrics were computed for all 19.264 genes and for the top 20 differentially expressed genes.

### Drug Response Prediction with Predicted Profiles

We implemented a sequential drug-response predictor that combines classification and regression. The logistic classifier used `threshold = 0.02` applied to `1 - AUDRC`. For regression, the ElasticNet model was trained on samples with `1 - AUDRC > 0.02`. Hyper-parameters (`alpha` and `l1_ratio`) were tuned using inner cross-validation. For samples classified as sensitive, the regression output was used; otherwise, predicted response was set to 0.

First, we trained drug response models on measured profiles of (cell line, drug) pairs in the training partition and evaluated them on either measured or predicted profiles belonging to pairs in the test partition. Next, we trained drug response models on profiles predicted by a PRM or baseline model, and evaluated this model on the profiles predicted by the same model.

### Gene Set Enrichment Analysis (GSEA)

Genes were ranked by the magnitude of their ElasticNet coefficients (from large positive to large negative). GSEA was performed using the gseapy package, utilizing the MSigDB Hallmark gene sets (Liberzon et al. [2015]) with 1,000 permutations. Normalized Enrichment Scores (NES) and False Discovery Rates (FDR) were employed to assess significance.

### Gene–Gene Correlations

We selected the 100 genes with the largest absolute average weights in the models trained on measured profiles. For each (cell line, drug) pair, correlations between gene expression and 1-AUDRC values were computed using the Pearson correlation coefficient. Similarity between vectors of measured and predicted correlations was calculated using the cosi ne similarity.

## Data Availability

All datasets used in this study are publicly available. The McFarland dataset was downloaded from https://figshare.com/s/139f64b495dea9d88c70 The Sciplex3 dataset was downloaded from https://www.ncbi.nlm.nih.gov/geo/query/acc.cgi?acc=GSE139944.

## Code Availability

The code to reproduce the figures will become available at: https://github.com/ NiekBrouwer98/drug-response-prediction-with-PRMs-paper

## Acknowledgements

We thank all the members of the Delft Bioinformatics Lab and NKI Wessels Lab for thoughtful discussions and feedback. We thank the developers of CPA, GEARS, and scFoundation for making their models publicly available. This research is part of the Oncode Accelerator Project that has received funding from the Dutch National Growth Fund (NGF).

## Author Contributions

N.B., D.J.V., M.J.T.R. and L.F.A.W. conceived the study. N.B. implemented the methods, performed the analyses, and created the figures. M.D. assisted with implementing CPA. J.A. assisted with implementing the GEARS variants. M.J.T.R. and L.F.A.W. supervised the study. N.B. and L.F.A.W. wrote the manuscript with input from all authors.

## Extended Data

**Extended Data Fig. 1:**
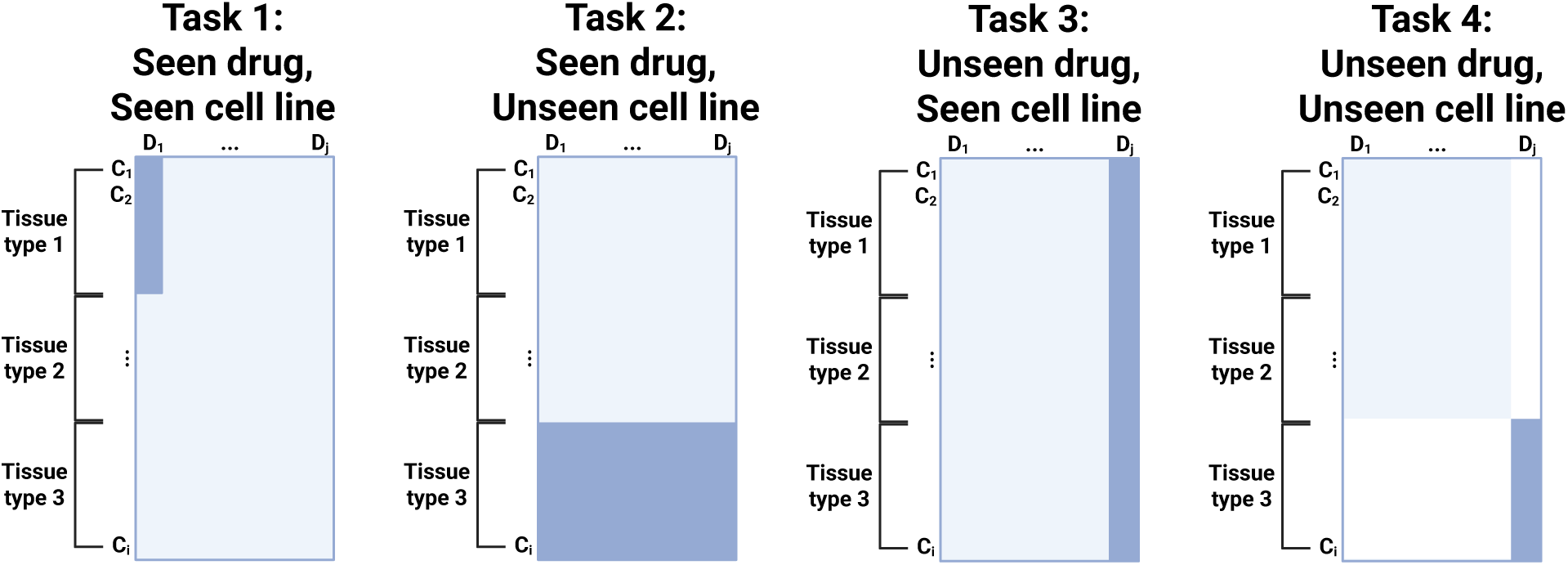
Overview of prediction tasks. Dark blue corresponds to the test set. To ensure that models do not leverage information from related cell lines, when a cell line is unseen, all cel lines from the same tissue type are also unseen. Models are evaluated with exhaustive leave-one-out cross-validation.

**Extended Data Fig. 2:**
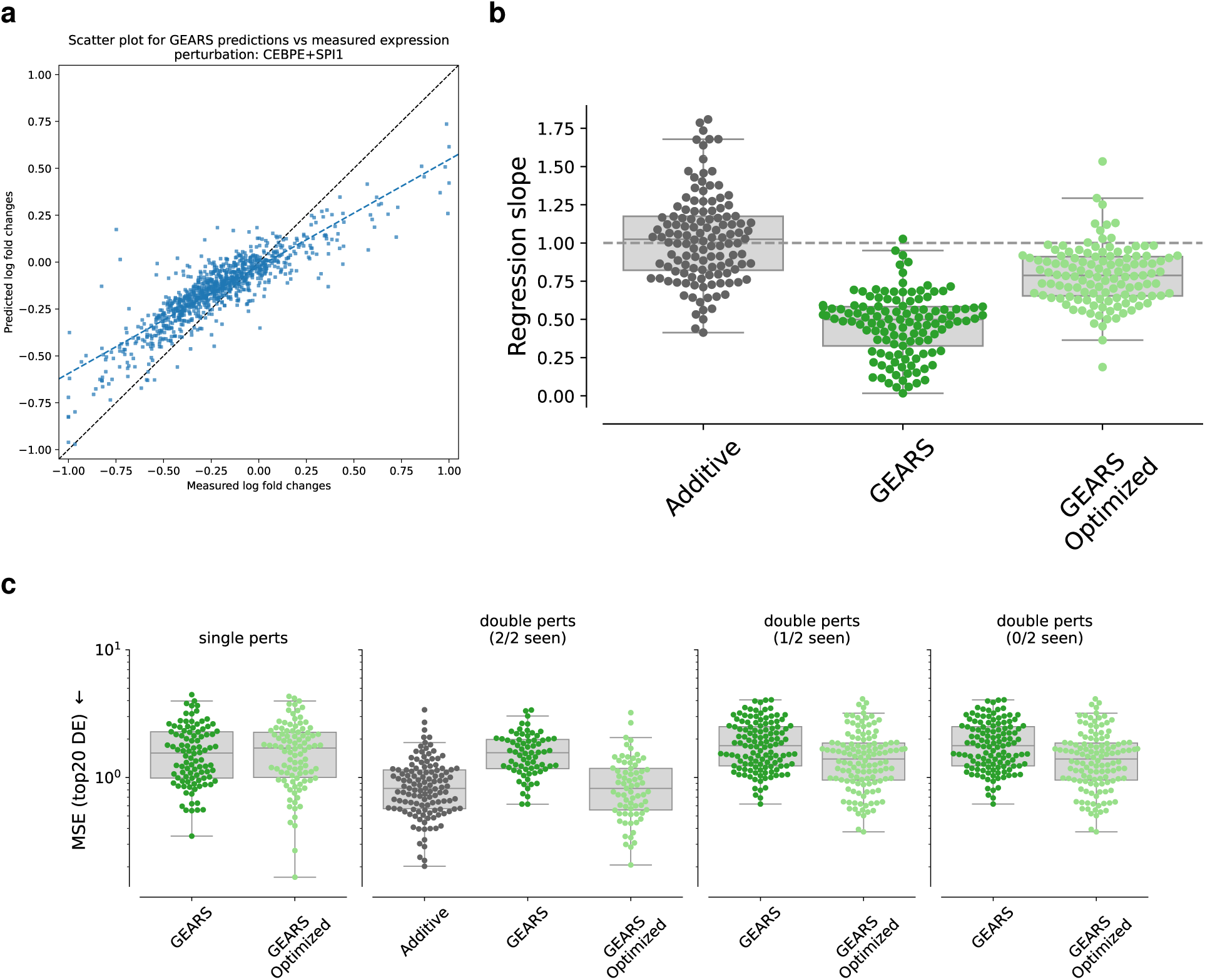
Performance overview of GEARS for Norman dataset (double-CRISPRa perturbation screen). **a,** Scatter plot of predicted vs. measured log fold changes for the double perturbation of CEBPE and SPI1 in the Norman dataset. Each point represents a gene, and the dotted line indicates the ideal y=x relationship. Values are consistently predicted closer to zero than the true values indicating underfitting of the model. **b,** Slopes of predicted vs. measured log fold changes for all double perturbations in the Norman dataset. Profiles are predicted with an additive model (sum of individual drug predictions), original GEARS model, and GEARS without regularization mechanisms (GEARS Optimized). Each boxplot summarizes the distribution of slopes for all genes across all double perturbations. **c,** Performance overview of additive baseline, GEARS original model, and GEARS optimized model for the Norman dataset across the top20 differentially expressed genes.

**Extended Data Fig. 3:**
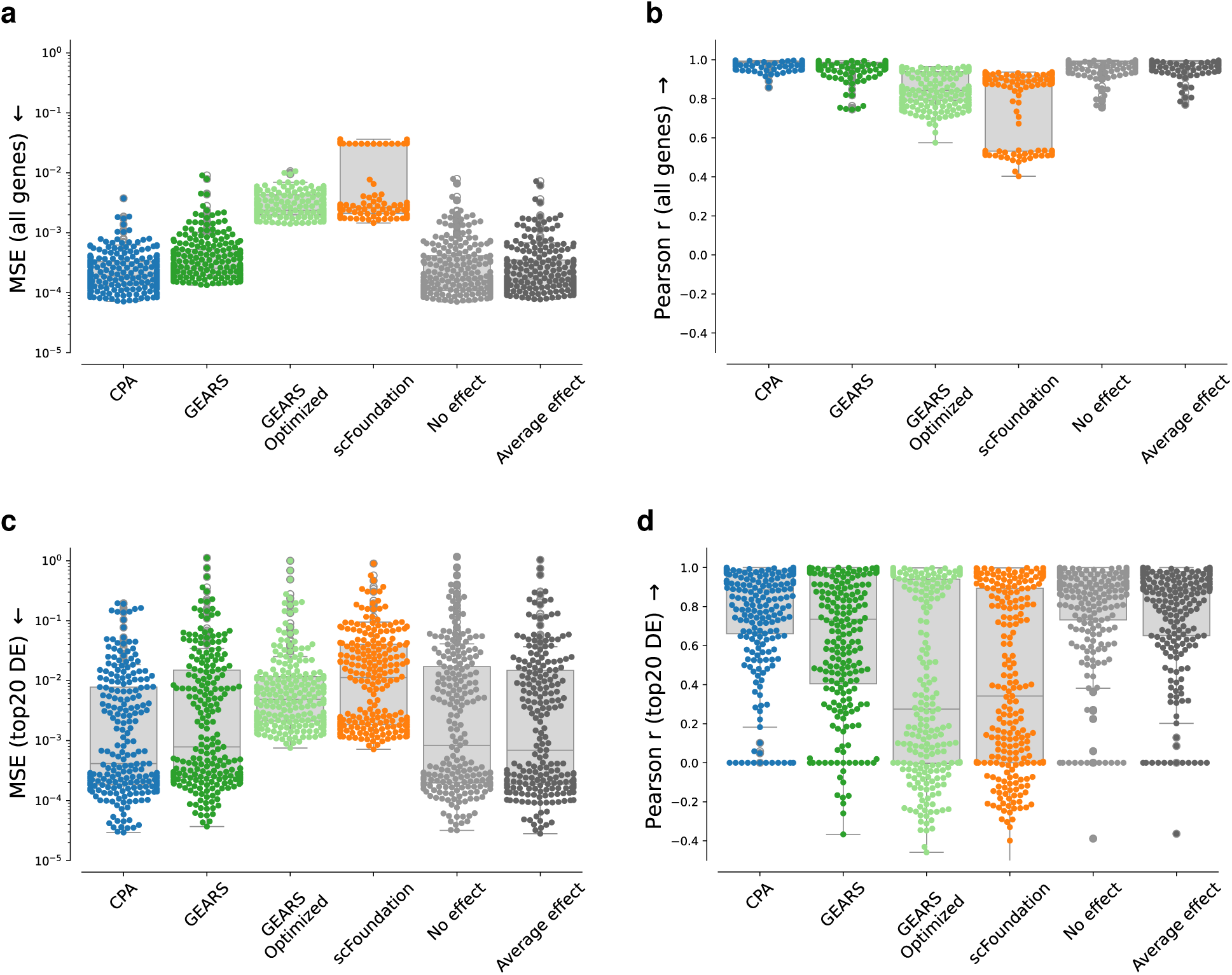
Performance overview of perturbation response models and baseline models with reconstruction accuracy metrics. **a,** Mean Squared Error (MSE) of predicted vs. measured post-treatment gene expression profiles across all genes. **b,** Pearson correlation coefficient (PCC) across all genes. **c,** MSE across top20 differentially expressed (DE) genes. **d,** PCC across top20 DE genes.

**Extended Data Fig. 4:**
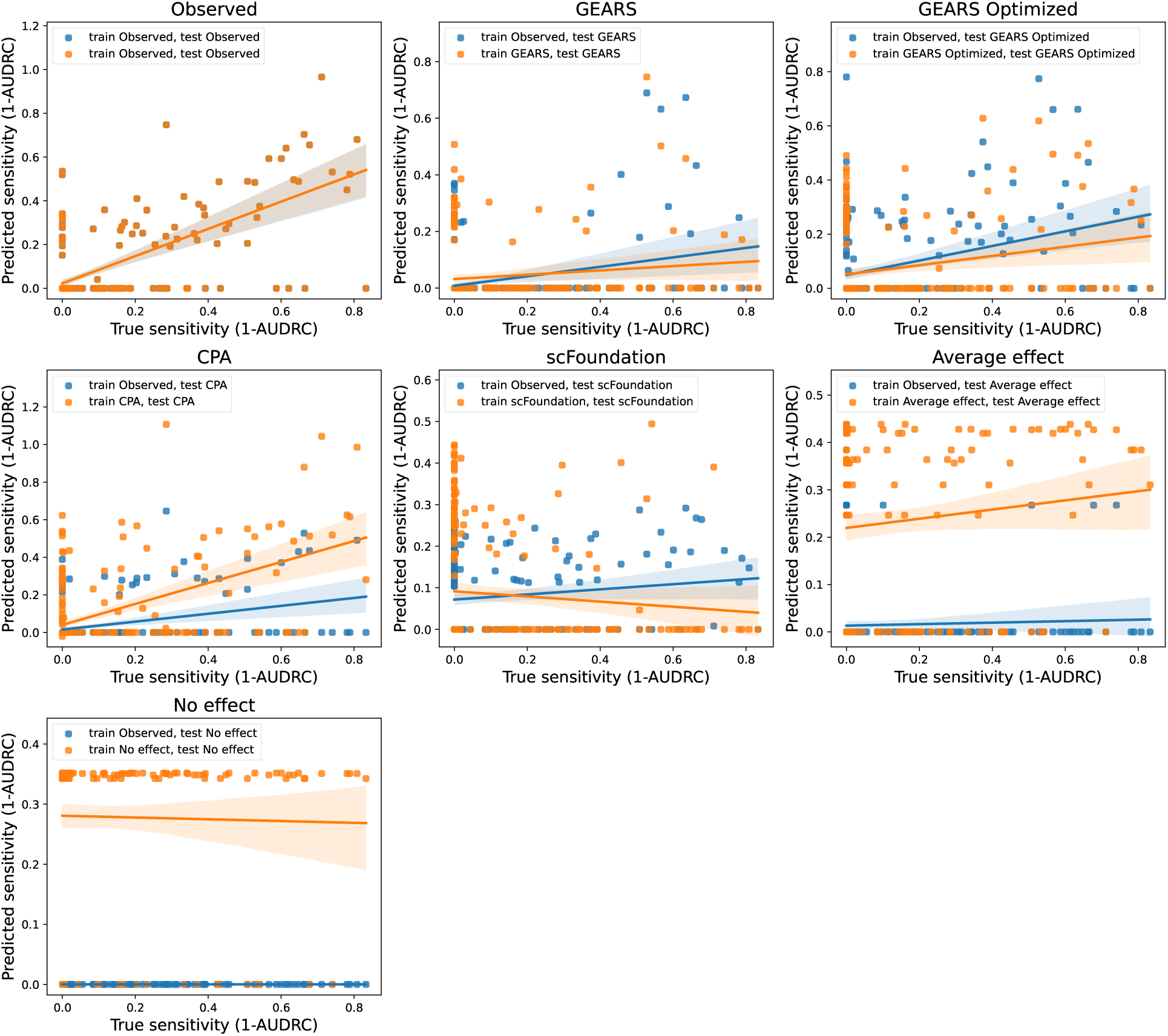
Drug response predictions. Drug response models are either trained on measured or predicted profiles. For each test split, performance is measured using Pearson correlation between predicted and observed AUDRC values.

**Extended Data Fig. 5:**
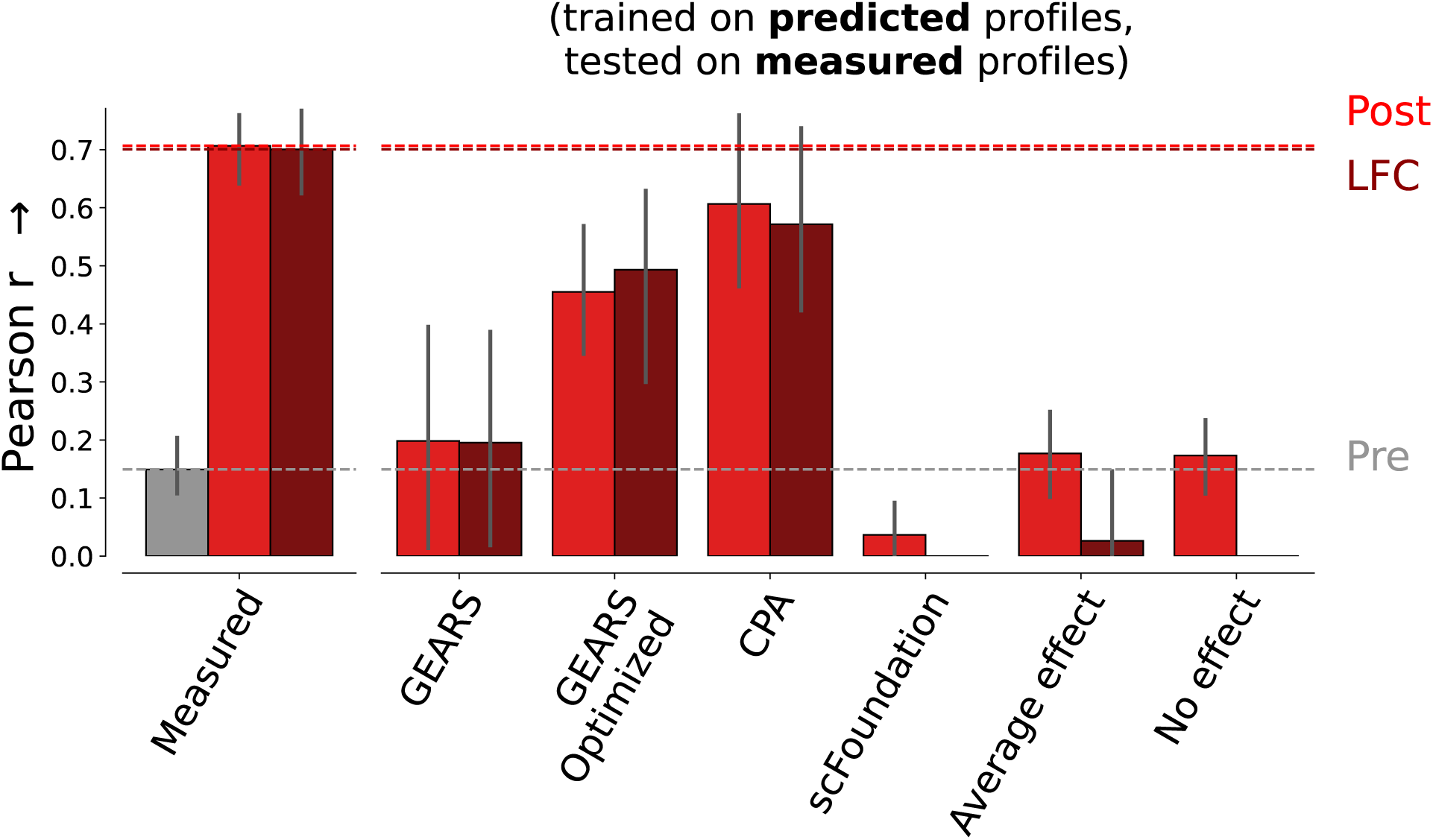
Performance overview of drug response predictions trained on predicted profiles and tested on measured profiles. For each test split, performance is measured using Pearson correlation between predicted and observed drug AUDRC values.

**Extended Data Fig. 6:**
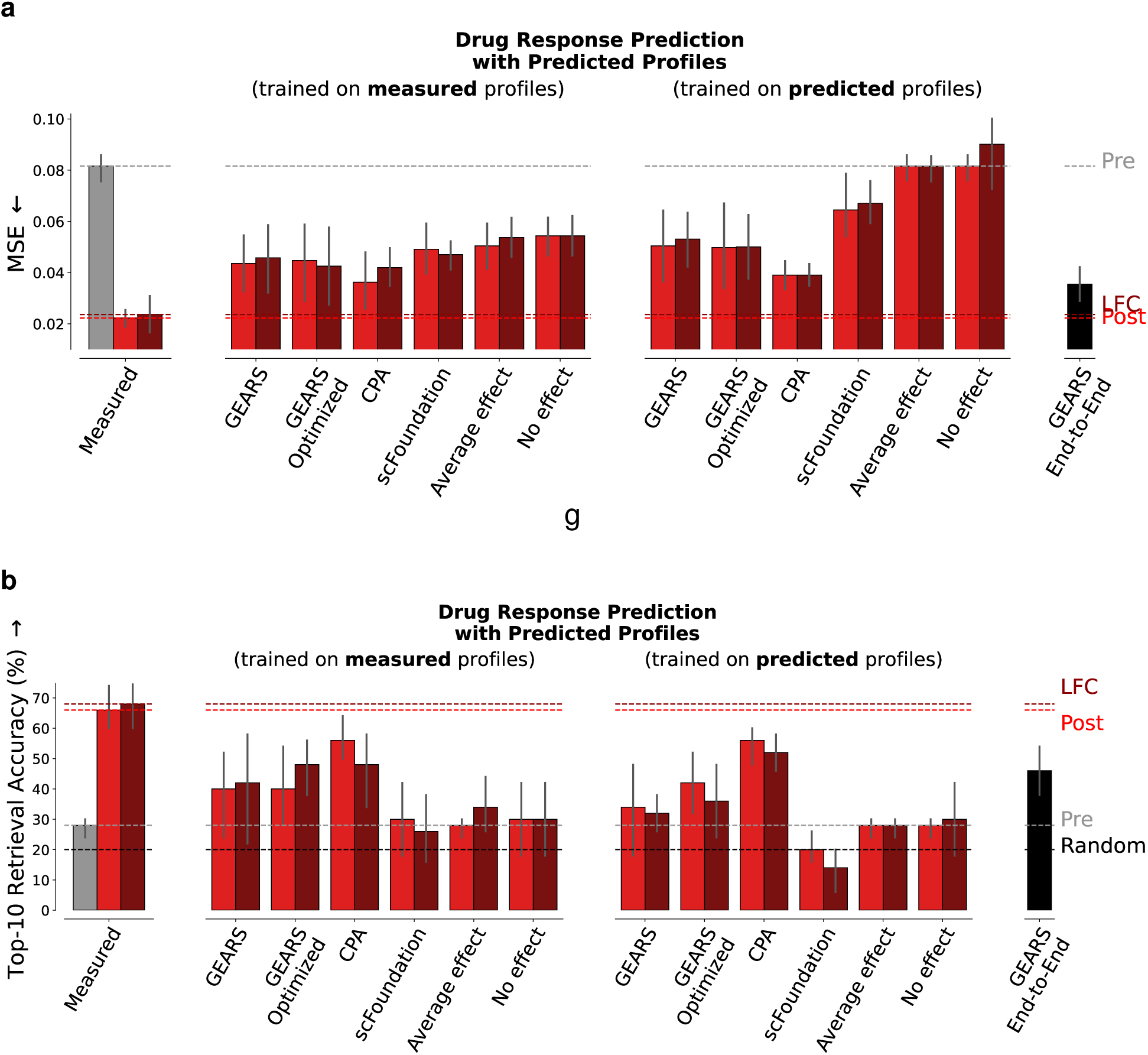
Performance overview of drug response prediction with alternative performance metrics. **a,** Evaluation using Mean Squared Error (MSE) of predicted and observed AUDRC values. **b,** Evaluation using retrieval accuracy (top 10% true sensitive (cell line, drug) pairs in top 10% predicted sensitive (cell line, drug) pairs).

**Extended Data Fig. 7:**
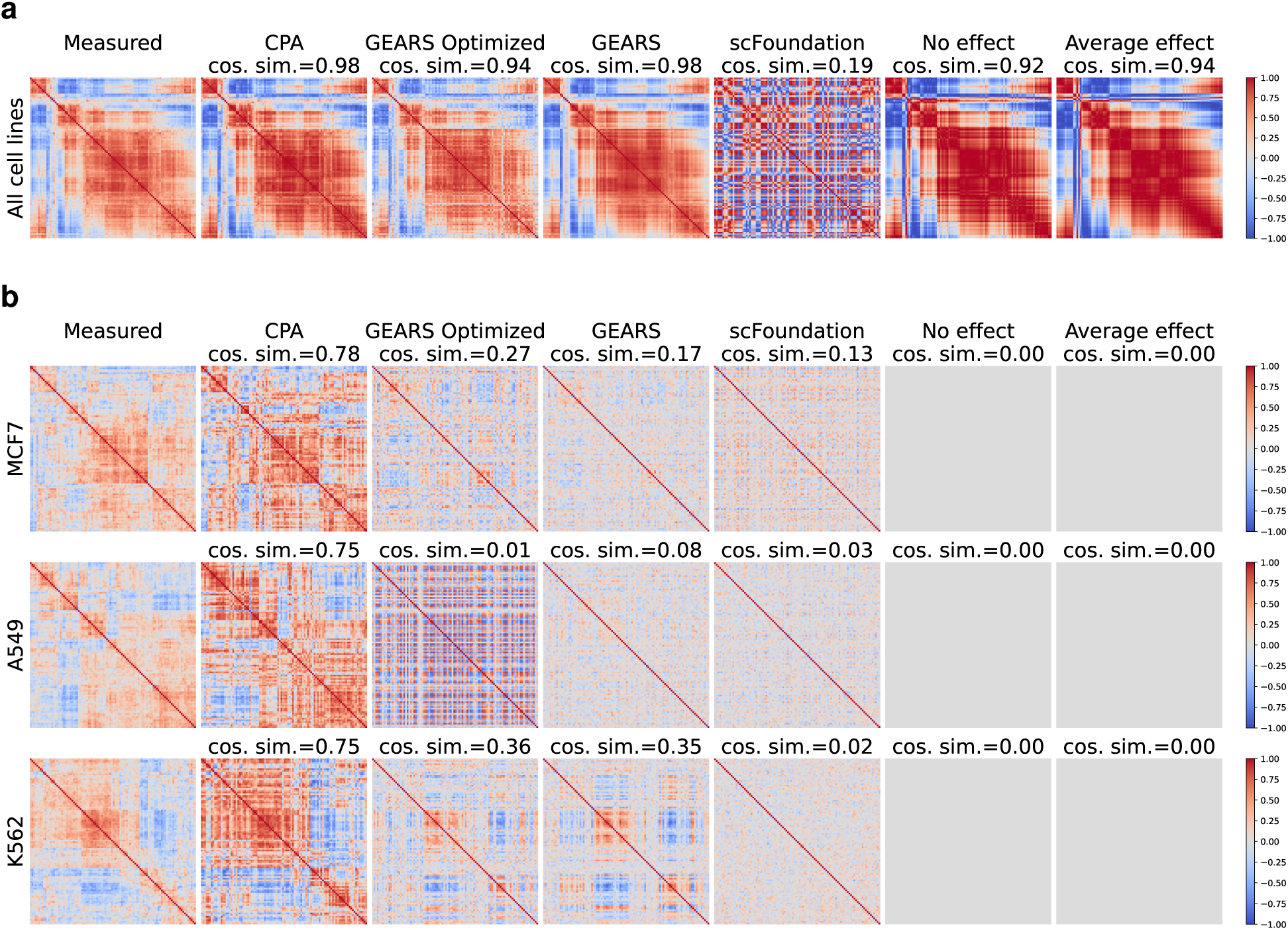
Gene-gene correlations in measured and predicted post-treatment gene expression profiles. **a,** Correlation structure of genes selected by drug response models trained on measured profiles. Correlations are computed across cell lines. **b,** Correlations computed per cell line. The correlation structure in CPA-predicted profiles is comparable to the measured profiles across all three cell lines.

**Extended Data Fig. 8:**
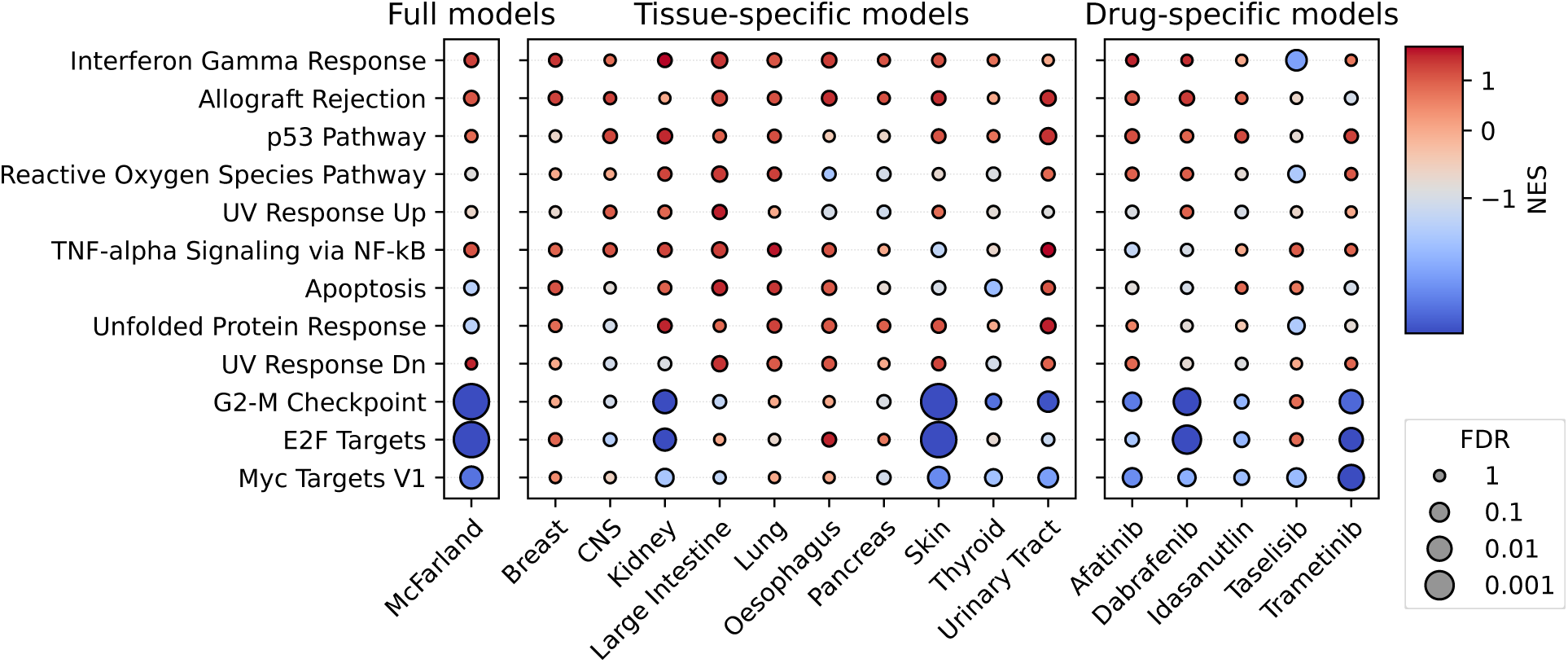
Gene set enrichment of genes selected by drug response models trained on measured gene expression profiles for the McFarland dataset. Models are either trained on the entire dataset, cell lines from the same tissue type (tissue-specific models) or cell lines treated with the same drug (drug-specific models). Dots are colored by Normalized Enrichment Score (NES), sized by adjusted p-value (FDR).

